# LINE-1 DNA methylation in response to aging and vitamin D

**DOI:** 10.1101/2020.05.02.073551

**Authors:** Lawrence T. C. Ong, Stephen D Schibeci, Nicole L Fewings, David R Booth, Grant P Parnell

## Abstract

Retrotransposons are genetic elements capable of their own propagation and insertion into the human genome. Because of their mutagenic potential, retrotransposons are heavily suppressed by mechanisms including DNA methylation. Increased age is associated with decreasing DNA methylation of the LINE-1 retrotransposon and may partially explain the predisposition towards malignancy with advancing age. Vitamin D has been investigated for its effects on DNA methylation at LINE-1 elements with mixed results. This study hypothesised that LINE-1 DNA methylation is altered by vitamin D exposure and age. Using whole genome bisulfite sequencing of adult and newborn haematopoietic progenitors, DNA methylation at LINE-1 elements was not found to vary between cells cultured with or without calcitriol, both in adults and newborns. In contrast, several LINE-1 regions were found to be differentially methylated between adults and children, but these were not uniformly hypermethylated in paediatric cells. The results of this study suggest that at least in haematopoietic cells, vitamin D does not appear to affect LINE-1 methylation.

## Introduction

Transposable elements encompass a diverse group of genetic elements that are capable of replicating and inserting copies of themselves into the genome. Retrotransposons form part of this family, capable of reinsertion through RNA intermediates. The family of retrotransposons includes long terminal repeat elements (LTRs), long interspersed nuclear elements (LINEs) and short interspersed nuclear elements (SINEs) (1). In humans, they form a large proportion of our genetic material, for example, LINE-1 retrotransposons and Alu elements comprise 17-25% and 11% of the human genome respectively (2, 3). Retrotransposons are generally thought to have deleterious effects due to their ability to induce malignancy through mutagenic insertions. Some inflammatory diseases are also thought to arise from transcription of retrotransposon elements that act as potent promoters for inflammatory genes (e.g. interferon-responsive genes in Aicardi Goutieres syndrome), or through triggering of innate immunity via pattern recognition receptors (e.g. MDA5, Toll-like receptors) (4, 5). Retrotransposons like LINE-1 also enable transposition of non-autonomous retroelements and the HERV-W endogenous retrovirus subfamily (6) previously linked to multiple sclerosis.

DNA methylation involves the covalent addition of a methyl group to a cytosine residue. In general, DNA methylation is associated with compacted chromatin and transcriptional repression. Retrotransposons are routinely methylated to decrease the risk of unwanted transposition and mutagenesis (7, 8). However, DNA methylation at these sites is also known to decrease with age (9–11), particularly at Alu elements (10), perhaps, in part, explaining susceptibility to malignancy with advancing age.

Vitamin D is a hormone that has pleiotropic effects, with important roles in physiologic homeostasis, disease pathogenesis and organism development. It is an important mediator of risk in autoimmune and allergic diseases such as multiple sclerosis (12), type 1 diabetes (13) and atopic dermatitis (14). In addition, it has been shown to have antiproliferative effects both in healthy and malignant cells (15). Several studies support the notion that the effects of vitamin D might be mediated, at least in part, through its effects on DNA methylation (see (16) for a review).

Studies examining the effect of vitamin D on retrotransposon DNA methylation have exclusively focused on LINE-1 DNA methylation. In humans, studies on peripheral blood have not found any changes in LINE-1 DNA methylation due to vitamin D. In one study, 50 individuals were supplemented with vitamin B and D, with no detectable changes in whole blood LINE-1 methylation after one year (17). Another study on peripheral blood lymphocytes, did not find any association between serum calcidiol and LINE-1 methylation (18). Studies on colorectal tissue have found higher serum vitamin D levels to be associated with increased LINE-1 methylation (19), perhaps explaining its effect on decreasing colorectal cancer risk. A study on visceral adipose tissue found a positive correlation between vitamin D levels and LINE-1 DNA methylation in colorectal cancer patients, but not controls (20).

In the present study, it was hypothesised that cell culture of haematopoietic progenitor cells with vitamin D, would result in changes in LINE-1 DNA methylation. Given the known effects of age on DNA methylation, it was also hypothesised that LINE-1 DNA methylation would be increased in cells derived from children compared to those from adults.

## Methods

### Cell isolation & culture

The methods have been previously described elsewhere (21). In brief, positive selection of CD34+ haematopoietic progenitors was performed on peripheral blood from two healthy adult male platelet donors (27 and 47 years old) and two umbilical cord blood units from one male and one female donor. Purified cells were plated at a density of 5×10^4^ cells/ml of media. The culture cocktail contained X-VIVO10 (Lonza), Albumex (Seqirus) 0.05%, SCF (Peprotech) 200ng/ml, GM-CSF (Peprotech) 0.03ug/ml, M-CSF (premium grade; Miltenyi Biotec) 5000U/ml, IL-6 (Peprotech) 10ng/ml, FLT3 ligand (Peprotech) 50ng/ml and Gentamicin (Sigma Aldrich) 50μg/ml. Calcitriol (1,25(OH)_2_Vitamin D_3_; BioGems) was added at a physiological concentration of 0.1nM. Cells were incubated at 37°C with 5% CO_2_ for one week before replating at a density of 1×10^5^ cells/ml of media. Media was then changed every third day by demi-depletion. Cells were harvested on day 21 and FACS sorting was performed to obtain purified CD14+CD45+ mononuclear phagocytes.

### Whole genome bisulfite sequencing, data QC, alignment and processing

Whole genome bisulfite sequencing libraries were generated with the Accel-NGS Methyl-seq DNA Library Kit (Swift Biosciences) and sequenced on a HiSeq X10 (Illumina) in 150bp PE mode with PhiX spike-in to counteract low sequence diversity. The quality of raw sequences was ascertained using FastQC (22). Quality trimming was carried out using Trim galore (23) in paired end mode with the following parameters -quality 20, --three_prime_clip_R1 5, --clip_R2 15. Reads were aligned to the hg19 genome using the Wildcard Alignment Tool (WALT) before further processing with Methpipe (24). Methylation calls were made using *methcounts* (using the -n option for CpG context cytosines only) before *symmetric-cpgs* was used to extract and merge methylation data from both strands. *Merge-methcounts* was used to merge locus specific read counts. Regional methylation analysis was performed using the *roimethstat* module (with -P and -v options), to determine methylation state within a prespecified region of interest. Sex chromosomes were omitted from analyses. Unless otherwise specified, analyses were performed on filtered data with an average read count of ≥ 10 reads per covered CpG for the entire region. Comparisons were made on regions where all samples met the filtering criteria. Global estimates of LINE-1 DNA methylation were determined by dividing methylated reads by total reads.

### Differential methylation and expression analysis

RADmeth (25) was utilised for differential methylation analysis. Unfiltered, CpG wise methylation counts from annotated LINE-1 regions were used as input. The -bins parameter was set to 1:200:1 and a CpG-wise FDR-adjusted significance of 0.05 was used to determine differentially methylated sites. The effects of calcitriol and age were considered by comparing the effects of calcitriol amongst cells of adult and paediatric origin separately. LINE-1 annotations were derived from Repeatmasker (26). The intersection of LINE-1 regions, annotated CD14+ promoter regions (27) and hg19 genes was determined with BEDtools (28). Details of the differential expression analysis can be found in (21).

## Results

On average, 96% of genome wide CpGs were covered at a depth of 16x. Approximately 65% of all annotated LINE-1 regions were covered (see Table 1). We had previously shown that across all genome wide CpGs, the average methylation was 0.83 irrespective of age or vitamin D. Unsurprisingly, LINE-1 regions demonstrated higher DNA methylation levels than those estimated at all genome wide CpGs and was slightly higher in paediatric than adult cells (0.89 vs 0.88; see Table 2 for descriptive statistics).

**Table 1.**
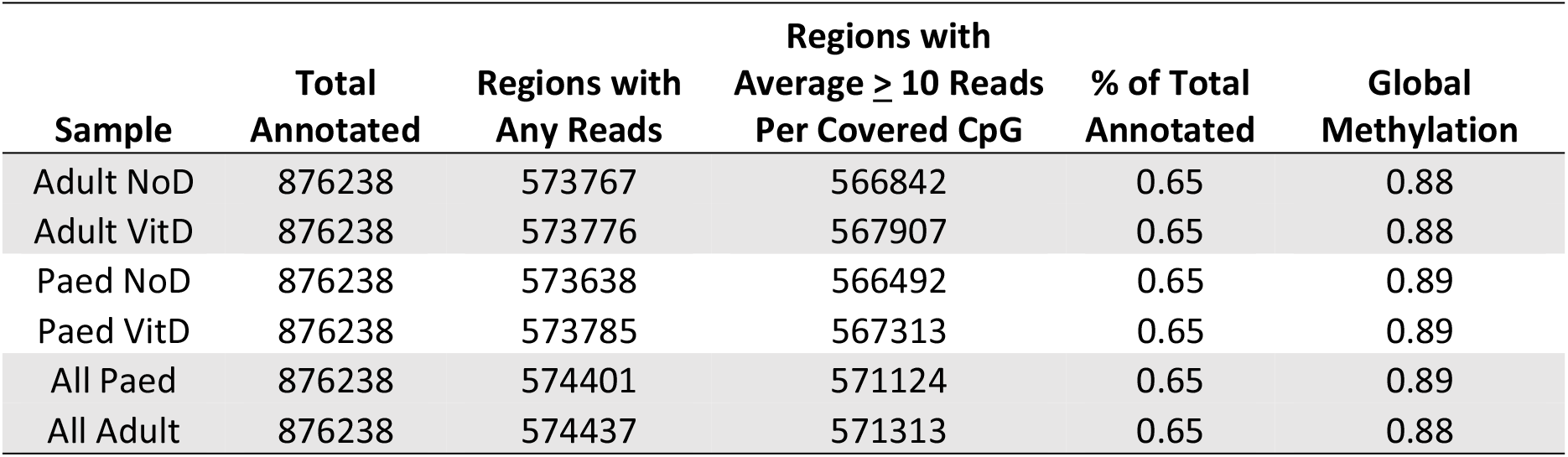
Coverage statistics of genome-wide annotated LINE-1 regions

**Table 2.** Descriptive statistics of DNA methylation for filtered LINE-1 regions

| Sample | Max | Min | Median | Mean | IQR |
| --- | --- | --- | --- | --- | --- |
| Adult NoD | 1 | 0 | 0.92 | 0.88 | 0.11 |
| Adult VitD | 1 | 0 | 0.91 | 0.88 | 0.11 |
| Paed NoD | 1 | 0 | 0.92 | 0.89 | 0.11 |
| Paed VitD | 1 | 0 | 0.93 | 0.89 | 0.10 |
| All Paed | 1 | 0 | 0.92 | 0.89 | 0.10 |
| All Adult | 1 | 0 | 0.91 | 0.88 | 0.11 |

IQR – interquartile range

There were no differentially methylated CpGs when comparing adult CD14+ with or without calcitriol (Figure 1 A & B). Similarly, no CpGs were found to be differentially methylated amongst paediatric CD14+ in the presence or absence of calcitriol (Figure 1 C & D). Interestingly, when comparing adult and paediatric data, there were 12450 differentially methylated LINE-1 CpGs corresponding to 5356 different LINE-1 regions (Supplemental Data 1 & 2, Figure 1 E & F). Of the differentially methylated CpGs, DNA methylation at 5160 CpGs was lower in cells of paediatric origin and higher in 7290.

**Figure 1.**
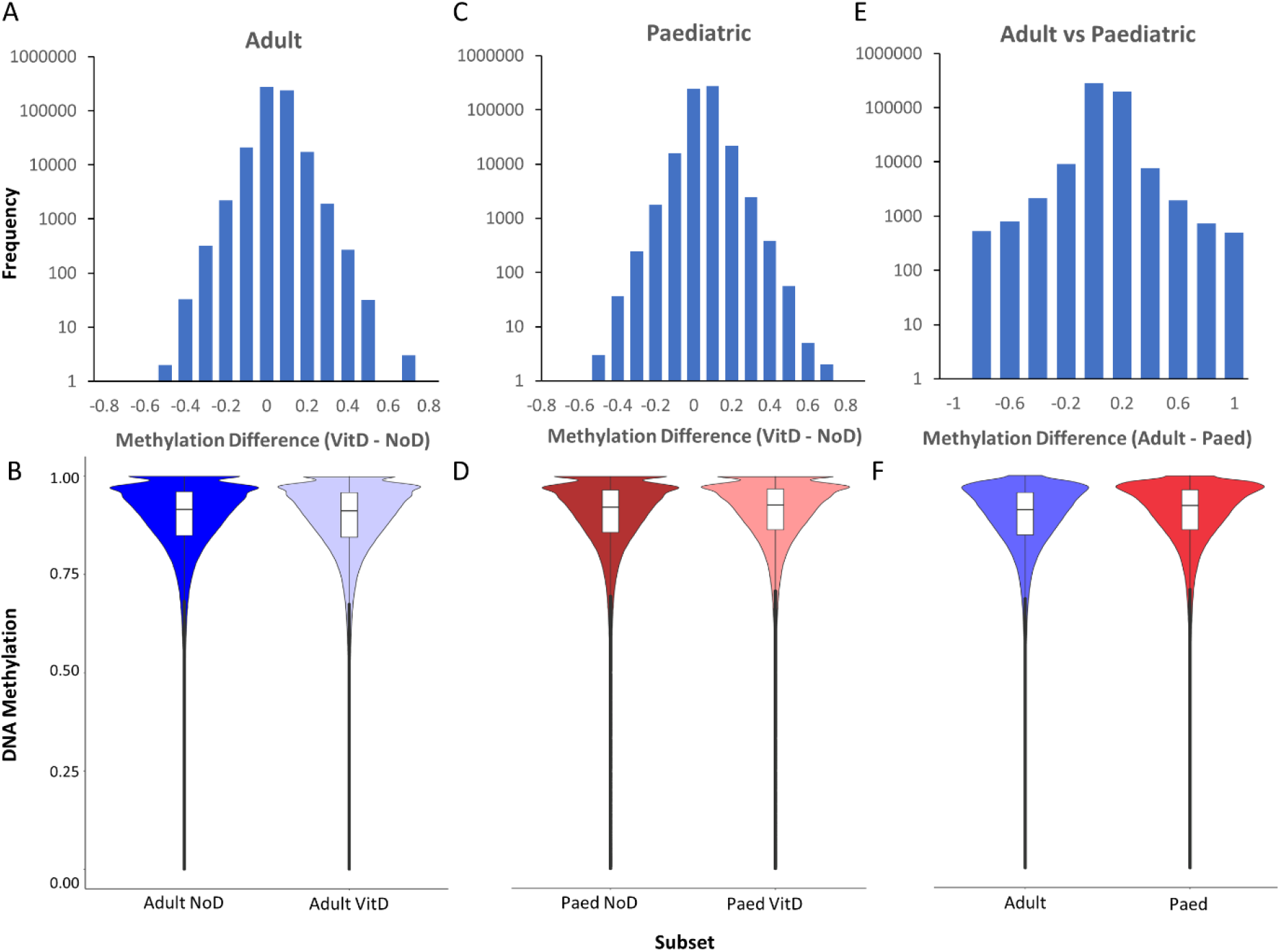
Histograms demonstrating DNA methylation differences between cells with and without calcitriol amongst **A)** adult cells and **C)** paediatric cells. **E)** compares all adult cells to paediatric cells. The lower plots, **B), D)** and **F)** display the distribution of DNA methylation values at genome wide LINE-1 regions corresponding to the above histograms. Paed – paediatric. NoD – without vitamin D. VitD – with vitamin D

Differentially methylated LINE-1 regions were further examined by determining their overlap with annotated CD14+ promoter regions and their corresponding hg19 genes. This yielded 312 genes (Supplemental Data 3) of which 46 were also differentially expressed (Supplemental Data 4). Several of these differentially methylated and differentially expressed genes have immunoregulatory functions.

## Discussion

Vitamin D has broad ranging physiological effects, of which a subset may be mediated through DNA methylation. This study explored the effects of vitamin D and age on LINE-1 DNA methylation. Consistent with previous studies on haematopoietic cells, there was no effect of vitamin D on LINE-1 methylation. In contrast, age appeared to be associated with changes in almost 11% of the interrogated LINE-1 regions. The differentially methylated CpGs varied with regard to the direction of differences. Although most loci were more highly methylated in cells of paediatric origin, a large proportion (41%) were less methylated. Thus, LINE-1 methylation is not only locus specific, but also variable with regard to the direction and magnitude of differences across age.

The results of this study suggest that vitamin D exposure during haematopoietic cell differentiation does not affect DNA methylation at LINE-1 sites. This was consistent with the two previously identified studies on haematopoietic cells. These results may be reconciled with those examining tissue LINE-1 methylation, by the fact that DNA methylation in different cell types appears to vary in its susceptibility to vitamin D induced alterations. These results suggest that, at least in myeloid cells, vitamin D may not act through LINE-1 DNA methylation to alter genomic stability or propensity to disease.

With respect to age related differences in LINE-1 DNA methylation, the overall magnitude of these differences at the global level was small. Previous studies examining age and LINE-1 methylation have studied adults. To date, the trajectory of global LINE-1 methylation loss has not been determined and this study suggests that changes may be smaller in childhood. Of particular interest however, was that the direction of differences varied between LINE-1 regions. This may point to the need for more targeted assessment of LINE-1 DNA methylation, rather than assuming that changes in their methylation are unidirectional across loci. Age related differences in LINE-1 DNA methylation also occurred at different gene promoters, a subset of which were associated with changes in gene expression. This may indicate a regulatory role of DNA methylation at some of these LINE-1 regions.

One of the strengths of this study was that we utilised a number of strategies to increase the likelihood of detecting DNA methylation changes. DNA methylation is known to be more plastic in early life (29), and susceptible to environmental stimuli during differentiation (30). The comparison of cells originating from adults and newborns, and differentiation of CD34+ haematopoietic progenitors allowed us to take advantage of these previous findings. Secondly, we examined a single subset of leukocytes, decreasing the likelihood of DNA methylation changes being obscured by heterogeneous subsets. Finally, by performing an *in vitro* study, the potential for confounding variables between groups was controlled for compared with the previously reported *ex vivo* studies.

Perhaps adversely influencing the results of this study were the use of cultured cells, whose DNA methylation has previously been shown to increase with passage number (31). This may have obscured differences in DNA methylation due to vitamin D. In addition, the use of these cultured cells may not reflect DNA methylation states in primary cells. Finally, whole-genome bisulfite sequencing captured only 65% of LINE-1 regions and may not be reflective of DNA methylation changes in the remaining 35% of LINE-1 regions. We speculate that the relatively low proportion of LINE-1 regions covered may be related to repetitive nature of LINE-1 sequences, making alignment difficult.

The results of this study reiterate the lack of vitamin D effects on LINE-1 methylation in haematopoietic cells and the likely differential effects of vitamin D in different tissues. Age related differences in LINE-1 DNA methylation were found, although the relationship is not as clear as that portrayed by global assessments of LINE-1 DNA methylation. Future studies might consider the rate of change in LINE-1 DNA methylation across the life span, and locus specific methylation to better elucidate the functional consequences of age related differences.

## Declarations

### Ethics approval and consent to participate

This study received ethics approval from the Western Sydney Local Health District Human Research Ethics Committee (HREC2002/9/3.6(1425) & (5366) AU RED LNR/17/WMEAD/447).

### Consent for publication

Not applicable

### Availability of data and materials

The datasets used and/or analysed during the current study are available from the corresponding author on reasonable request.

### Authors’ Contributions

LO, GP and DB devised the experiments. NF and SS assisted in planning and analysis of cell culture and flow cytometric experiments. LO conducted the experiments, analyses and prepared the manuscript. GP performed RNA-seq and assisted in data analysis. All authors read and approved the final manuscript.

## Acknowledgements

The authors would like to acknowledge Australian Red Cross Blood Services, the Sydney Cord Blood Bank and donors for providing samples for this study. Flow cytometry was performed at the Flow Cytometry Core Facility supported by the Westmead Research Hub, Cancer Institute NSW and NHMRC. Bioinformatic analysis was supported by Sydney Informatics Hub, funded by the University of Sydney.

## Supplemental Data

**Supplemental Data 1 –** Differentially methylated LINE-1 CpGs

**Supplemental Data 2 –** LINE-1 regions containing differentially methylated CpGs

**Supplemental Data 3 –** Genes corresponding to differentially methylated LINE-1 promoters

**Supplemental Data 4 –** Differentially expressed genes corresponding to differentially methylated LINE-1 promoters

